# Genomics and synthetic community experiments uncover the key metabolic roles of acetic acid bacteria in sourdough starter microbiomes

**DOI:** 10.1101/2024.04.11.589037

**Authors:** H. B. Rappaport, Nimshika P. J. Senewiratne, Sarah K. Lucas, Benjamin E. Wolfe, Angela M. Oliverio

**Affiliations:** Department of Biology, Syracuse University, Syracuse, NY, USA; Department of Biology, Tufts University, Medford, MA, USA

**Keywords:** acetic acid bacteria, sourdough starter, synthetic communities, comparative genomics, strain diversity, microbiome

## Abstract

While research on the sourdough microbiome has primarily focused on lactic acid bacteria (LAB) and yeast, recent studies have found that acetic acid bacteria (AAB) are also common members. However, the ecology, genomic diversity, and functional contributions of AAB in sourdough remain unknown. To address this gap, we sequenced 29 AAB genomes, including three that represent putatively novel species, from a collection of over 500 sourdough starters surveyed globally from community scientists. We found variation in metabolic traits related to carbohydrate utilization, nitrogen metabolism, and alcohol production, as well as in genes related to mobile elements and defense mechanisms. Sourdough AAB genomes did not cluster when compared to AAB isolated from other environments, although a subset of gene functions were enriched in sourdough isolates. The lack of a sourdough-specific genomic cluster may reflect a nomadic lifestyle of AAB. To assess the consequences of AAB on the emergent function of sourdough starter microbiomes, we constructed synthetic starter microbiomes, varying only the AAB strain included. All AAB strains increased acidification of synthetic sourdough starters relative to yeast and LAB by 18.5% on average. Different strains of AAB had distinct effects on the profile of synthetic starter volatiles. Taken together, our results begin to define the key ways in which AAB shape emergent properties of starters and suggest that differences in gene content resulting from intraspecies diversification can have community-wide consequences on emergent function.

**Importance:** This study is the first comprehensive genomic and ecological study of acetic acid bacteria isolated from sourdough starters. By combining comparative genomics with manipulative experiments using synthetic microbiomes, we demonstrate that even strains with >97% ANI can shift important microbiome functions, underscoring the importance of species and strain diversity in microbial systems. We also show the utility of sourdough starters as a model system to understand the consequences of genomic diversity at the strain and species level on multispecies communities. These results are also relevant to industrial and home-bakers as we uncover the importance of AAB in shaping properties of sourdough starters that have direct impact on sensory notes and quality of sourdough bread.

## Introduction

Acetic acid bacteria (AAB) are an important member of many fermented food microbiomes. They are part of the *Alphaproteobacteria* class, *Rhodospirillales* order, and *Acetobacteraceae* family, with over 100 species across 19 genera now described (1, 2). This group is well-recognized for the fermentation of vinegar (*A. pastuerianus* (1)), kombucha (*Komagataeibacter* spp. (3–5)), lambic beer (*A. lambici* (6)), water kefir (*Acetobacter sicerae* (7)), and cocoa (*A. pasteurianus* commercially, *A. ghanensis/senegalensis* spontaneously (8)), among others. Broadly, AAB produce acidic flavor in lambic beer and vinegar, contribute to flavor and discourage germination in cocoa beans, and produce the cellulose component of “SCOBY” in kombucha (7). AAB also release a variety of metabolic products that have been applied to food, cosmetics, medicine, and other industries (9). These products, including acetic acid (sour flavor and antimicrobial), DHA (common sunscreen ingredient), and acetoin (butter-like flavor) result from incomplete oxidative fermentation of sugars and alcohols (2, 10). AAB are also found in insect guts, including fruit flies and bees (11), as well as flowers and fruits (2), which are all natural sugary environments (12). Adaptation to these sugary environments may have facilitated success in fermented foods and beverages (7).

Despite their well-recognized importance, there is a limited understanding of the ecology of AAB. One system where AAB are likely important but have largely been overlooked is sourdough. The bulk of sourdough microbiome research has focused on yeast and lactic acid bacteria (LAB). These two functional groups are sufficient to make a sourdough starter (13), although starter microbiomes are often more diverse (with ∼3-10 species) and include other microbes beyond LAB and yeast (14–17). The “back slopping” method involved in the maturation and maintenance of sourdough starters allows for low abundance species to become more dominant and for new species to colonize, but factors that allow additional groups to persist are unclear (18). For example, recent work has indicated that AAB are commonly found in many sourdough starters (14, 19–21), but the importance and function of AAB within sourdough remains uncertain as well.

A few studies have begun to investigate the functional roles of AAB in sourdough, but genomic and metabolic characterizations of AAB have been limited to other environments including insect guts (11) and vinegar (22). Common garden experiments with wild sourdough starters suggest that AAB may affect dough rise and aroma (14), but there is limited controlled experimental validation. A strain of *A. tropicalis* is known to influence resultant properties of Chinese steamed bread (23, 24). Li et al. found that starters with AAB added had the lowest pH, highest viscosity and elasticity, and had a greater variety of flavor compounds than bread with yeast and LAB or yeast alone. Likewise, the addition of extracted exopolysaccharides (levans, fructans) from AAB to dough resulted in bread that was softer and had more volume, further indicating that the compounds released by AAB can impact emergent traits of sourdough (25). However, insights from the vast majority of ecologically and functionally diverse AAB remain limited.

In addition to their economic and cultural significance, AAB in sourdough also present an opportunity to study the ecological consequences of genomic and strain variation within microbiomes. While it is clear that intraspecies diversity exists across many microbiomes from comparative genomics and metagenomics studies (26–30), there are still surprisingly few studies that have experimentally manipulated strain diversity to understand impacts on microbiome composition and function. Past studies investigating the role of variation at the intraspecies level have found that strain-level differences are associated with microbial community composition and functional traits (31–33), and that multiple strains can coexist in an environment due to multiple-niche polymorphism at a small scale (34, 35). As members of microbiomes across a wide range of environments from sourdough to insects, AAB may be generalists, but environmental selection pressures may have resulted in intraspecies diversification. It is also unclear how much strain diversity exists within AAB species in sourdough and other environments and how this diversity contributes to variation in emergent functions of microbiomes. Understanding the functional significance of species and strain diversity within AAB may help reveal novel approaches for managing the functions of fermentations and other AAB-dominated microbiomes.

To shed light on the ecological and functional roles of AAB at multiple levels of genetic similarity in sourdough starter microbiomes, we isolated dominant AAB taxa from a diverse collection of 500 sourdough starters contributed by community scientists from around the world (14). This collection represents 21 strains across 11 species spanning two genera (**Fig. 1A**). We obtained high quality draft genomes of all isolates, and also obtained eight metagenome assembled genomes (MAGs), to characterize metabolic pathways and differences in gene content. We assessed if particular functions were enriched in sourdough AAB genomes versus those from other environments broadly across all AAB and within species clusters. We also used our sourdough AAB isolate collection to experimentally determine the function of diverse AAB within the sourdough starter microbiome. Constructing synthetic starter communities with and without various AAB strains, we measured key emergent properties such as acidification and metabolite production. Our work experimentally determines the consequences of an overlooked but functionally important group of microbes in sourdough starters, the acetic acid bacteria, on emergent microbiome function.

**Fig. 1:**
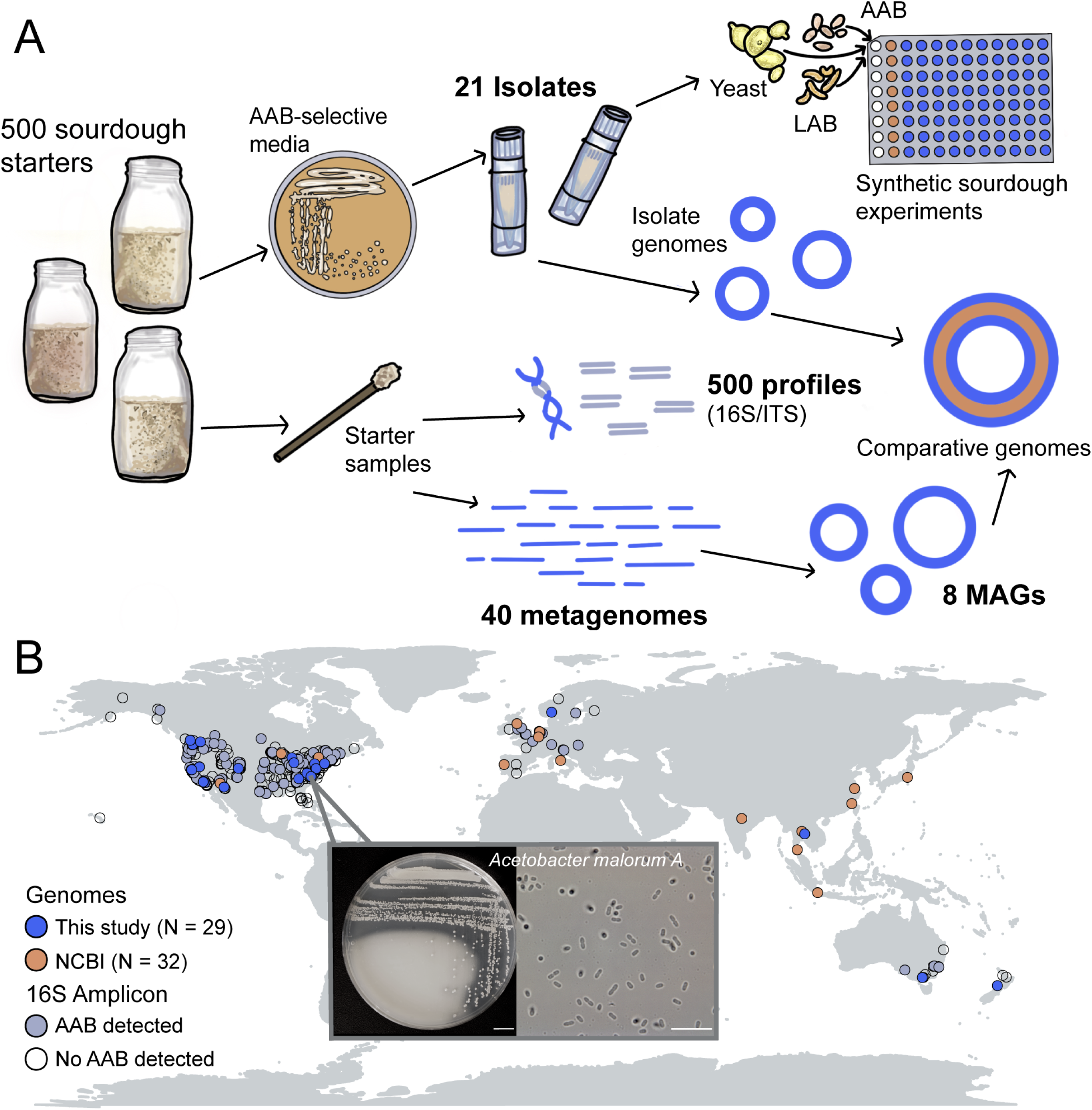
Overview of study design and geographic diversity of acetic acid bacteria. (A) Conceptual overview of study data and experiments. Leveraging 16S amplicon data from 500 sourdough starters, we isolated 21 AAB on selective media and obtained corresponding genomes. We obtained MAGs (N = 8) from metagenomes (sample N = 40) and also included 32 publicly available AAB genomes for a final genome set of 61 for our comparative genomics assessment. We also selected a subset of AAB isolates (N = 10) for synthetic sourdough experiments to measure the functional impact of AAB on starter microbiomes. (B) AAB were isolated from a set of 500 sourdough samples that were previously described using amplicon sequencing, collected from a global network of community scientists (14). Light blue gray circles indicate detection of AAB in 16S rRNA gene amplicon data and bright blue circles denote recovery of one or more AAB genomes from the sample. Orange circles denote the set of AAB genomes included from NCBI, recovered from diverse environmental sources and geographic locations. Popout shows example AAB colonies on plate (s.b. 1 cm) and under microscope at 100x (s.b. 10 µm).

## Results and Discussion

### Acetic acid bacteria are phylogenetically diverse and abundant in sourdough starter microbiomes globally

To determine the ecological distribution and diversity of AAB in sourdough starters, we leveraged a sample collection of 500 sourdough starters collected from a global network of community scientists (14). First, we did a detailed investigation of AAB across all starters (N=500) using a previously sequenced 16S rRNA amplicon dataset (14). Then, we obtained AAB genomes from the same starter collection by sequencing isolates and reconstructing microbial genomes from metagenomes. Across the 500 sourdough starters, AAB were present in ∼30% of samples (at ≥1% relative abundance). AAB comprised 23% of the overall bacterial community on average and up to 80% of the starter bacteria. We detected 26 AAB ASVs that span three genera (*Acetobacter*, *Gluconobacter*, *Komagataeibacter*, **Fig. S1**). In samples where AAB were present, 1.5 AAB ASVs were detected on average (**Table S1**).

We next investigated if AAB exhibited consistent patterns of co-occurrence with LAB, yeast, or other AAB, as this might help explain the variable distribution patterns of AAB across starters. We found that LAB including *Schleiferilactobacillus harbinensis*, *Lentilactobacillus kefiri,* and *Furfurilactobacillus rossiae* (**Table S2**) were enriched (FDR *P* < 0.001 for all) in AAB dominant samples (samples with >25% AAB). *S. harbinensis* and *L. kefiri* are common members of water kefir, an acidic beverage which is often populated by AAB, suggesting shared acid tolerance (36, 37). However, the most dominant LAB in sourdough (*Levilactobacillus brevis*, *Lactiplantibacillus plantarum*, *Pediococcus parvulus, Fructilactobacillus sanfranciscensis*) are found similarly with and without AAB. *Pichia mandshurica* was the only yeast significantly associated with AAB dominant samples. This species is associated with wine spoilage and produces acetic acid as well (38), likely explaining its success in AAB dominant sourdough samples with higher acidity. *Saccharomyces cerevisiae* made up about 75% of the yeast in both AAB dominant and non-AAB dominant samples, supporting that AAB can likely survive in most sourdough microbiomes. Within AAB, *Komagataeibacter* and *A. pasteurianus* were frequently detected in the same subset of samples (rho = 0.32, *P* < 0.001), indicating that multiple AAB can co-persist in starters (**Table S3**).

Leveraging our 16S amplicon data to guide culturing efforts, we isolated 21 AAB strains from 20 sourdough samples. The isolates we obtained span most of the geographic diversity of AAB from our sourdough starter collection (**Fig. 1B**). AAB were found in starters from 14 countries and four continents including the USA, Belgium, China, Thailand, New Zealand, Norway, and India. Reflecting our broader collection, AAB isolates lack representation from South America and Africa, and are overrepresented in North America, where we collected the most samples overall. We also added eight MAGs of AAB in sourdough from a set of 40 metagenomes **(Table S4**). We did not find any published AAB genomes isolated from sourdough starters based on our search of publicly available databases. Our AAB genomes represent a novel collection from sourdough starters. Notably, three of our recovered isolates likely represent putative novel AAB species based on <95% ANI match to described AAB species, distinct phylogenetic placement, and reported RED values from GTDB-Tk (39, 40). We anticipate the number of novel AAB lineages to continue to increase as AAB are further studied in sourdough and other previously overlooked fermentation environments (**Table S4**).

The AAB genomes we obtained represent most of the known phylogenetic diversity of AAB in sourdough starters (**Fig. 2**). Across the 500 starters, ASVs were assigned to 20 major clusters. Notably, genomes with as low as 90% ANI were assigned to the same ASV, despite representing multiple species, highlighting the limited ability of short-read amplicon sequencing to resolve genetic diversity of AAB. We obtained genomic representatives from eight of the ten most dominant clusters, as well as two representatives from less abundant clusters (**Fig. 2A**). The most abundant clusters in the 500 sourdough starters were *A. malorum*/*cerevisiae* and *A. oryzoeni*/*oryzifermentans*/*pasteurianus* at around 6% mean relative abundance. Our isolates reflect the dominant sourdough strains including: *A. malorum*, *A. pasteurianus*, *A. orientalis*, and *G. oxydans,* among others (**Fig. 2A**). Although we cultured most of the abundant groups, our efforts did not yield *Komagataeibacter* isolates.

**Fig. 2:**
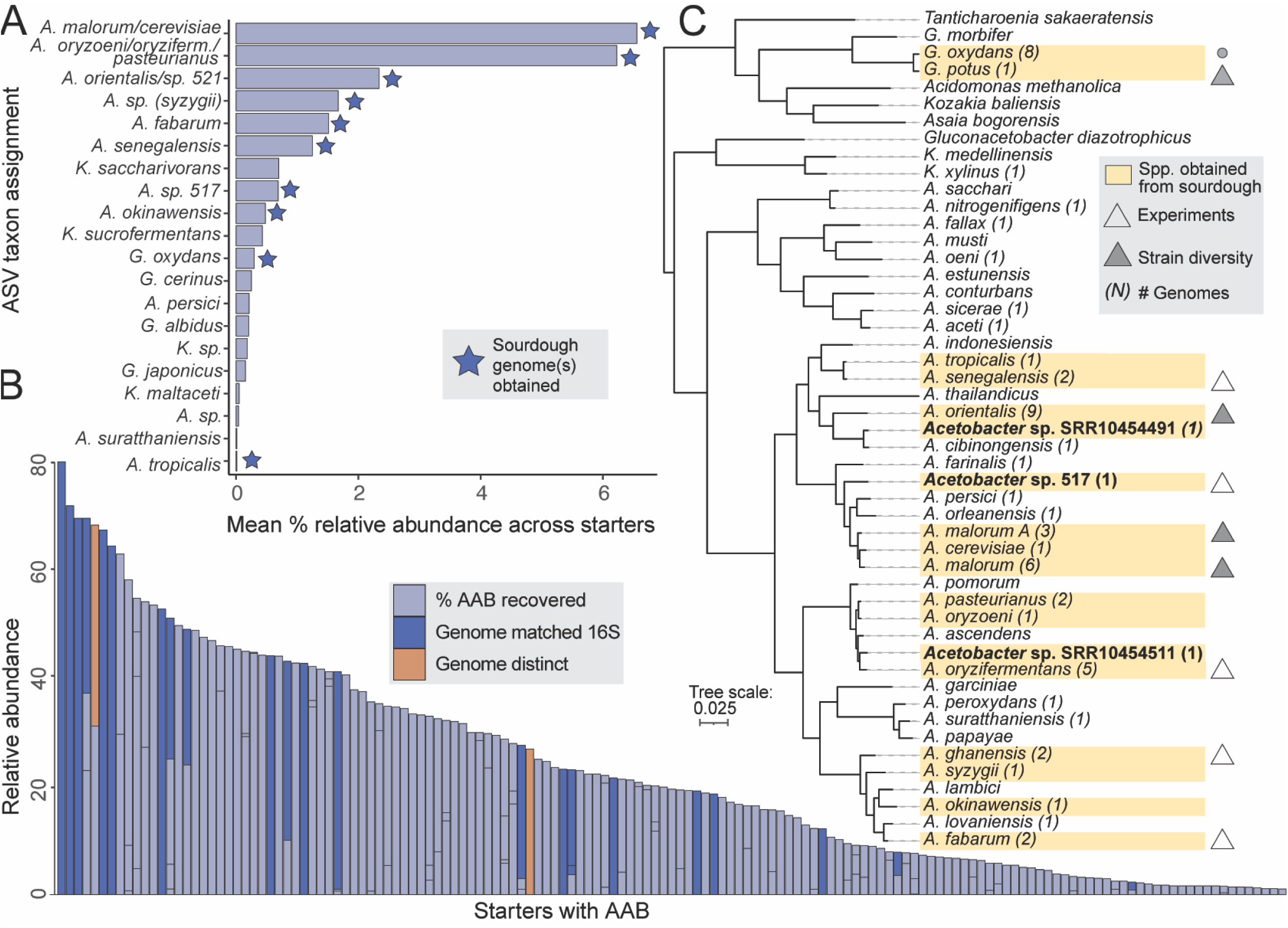
Genomic diversity and abundance of AAB in sourdough starter microbiomes. (A) Summary of the mean % relative abundance of AAB by species taxonomic assignments across 500 sourdough starters. Some ASVs could only be assigned to a nearest cluster of species (**Fig. S1**) due to low resolution of ASVs. Stars indicate species clusters for which we obtained one or more genomes. **(B)** Summary of the mean % relative abundance of AAB across starter samples. AAB were detected in 30% of the 500 starter samples at ≥1%. Breaks in bars represent the relative abundance of each AAB ASV detected in a sample. Dark blue bars denote samples where isolated taxonomy matched expected identity from 16S amplicon sequencing of community and orange denotes samples where a distinct AAB taxon was recovered. **(C)** Representative genome tree of *Acetobacter* species including three putatively novel species (in bold) and other genera including *Gluconobacter* and *Komagataeibacter*, which were also detected in sourdough starter microbiomes. The number of genomes obtained for this study are listed in parentheses. Species with isolates from sourdough or MAGs recovered from sourdough are highlighted in yellow. Triangles represent species that were included in our synthetic sourdough experiments, and filled in triangles are species that also were included in our assessment of intraspecies strain diversity.

There was high correspondence in the taxonomic identity of isolates obtained from our culturing efforts relative to expected AAB from amplicon sequencing (**Fig. 2B**), indicating that amplicon-informed targeted culturing is a tractable approach in this system and likely other fermented food systems where most members are readily culturable (41). The sourdough AAB genomes we obtained span much of the broader AAB diversity across environments as well (**Fig. 2C**), although a couple clades are notable as they were not detected across our 500 starters. For example, the clade that includes species such as *A. aceti, A. nitrogenifigens*, and *A. sacchari* does not contain any members detected in ASVs or isolated from sourdough.

### There is extensive variability in gene content and metabolic traits among sourdough AAB

To profile variation in gene content and metabolic traits of AAB genomes, we compiled a set of AAB genomes from sourdough and other environments. In addition to our 21 isolate genomes and eight MAGs, we included 40 high-quality AAB genomes from NCBI isolated from fermented beverages (beer, kombucha, cider), fruit, fruit flies, and a range of other sources (mud, tree bark, and sewage) resulting in a collection of 61 AAB genomes. NCBI genomes included three species we recovered from sourdough (*A. malorum*, *A. orientalis*, and *G. oxydans*) isolated from other sources, and other species of AAB (*A. fallax*, *A. lovaniensis*, *A. aceti*, and *K. xylinus*). Across AAB, differences in ANI ranged from 70% (across genera), a noted ANI gap of 90-95% observed in closely related species clusters, and 96-98% within species (**Table S5**). Across all genomes (N=61), we detected 11,596 unique gene clusters. Of these, 909 represented core gene clusters that were present in >95% of genomes (7.8%). The other 10,687 gene clusters were only detected in a subset of genomes and represent both species-specific clusters and other accessory genes that were variably detected (**Fig. 3A**, **Table S6**).

**Fig. 3:**
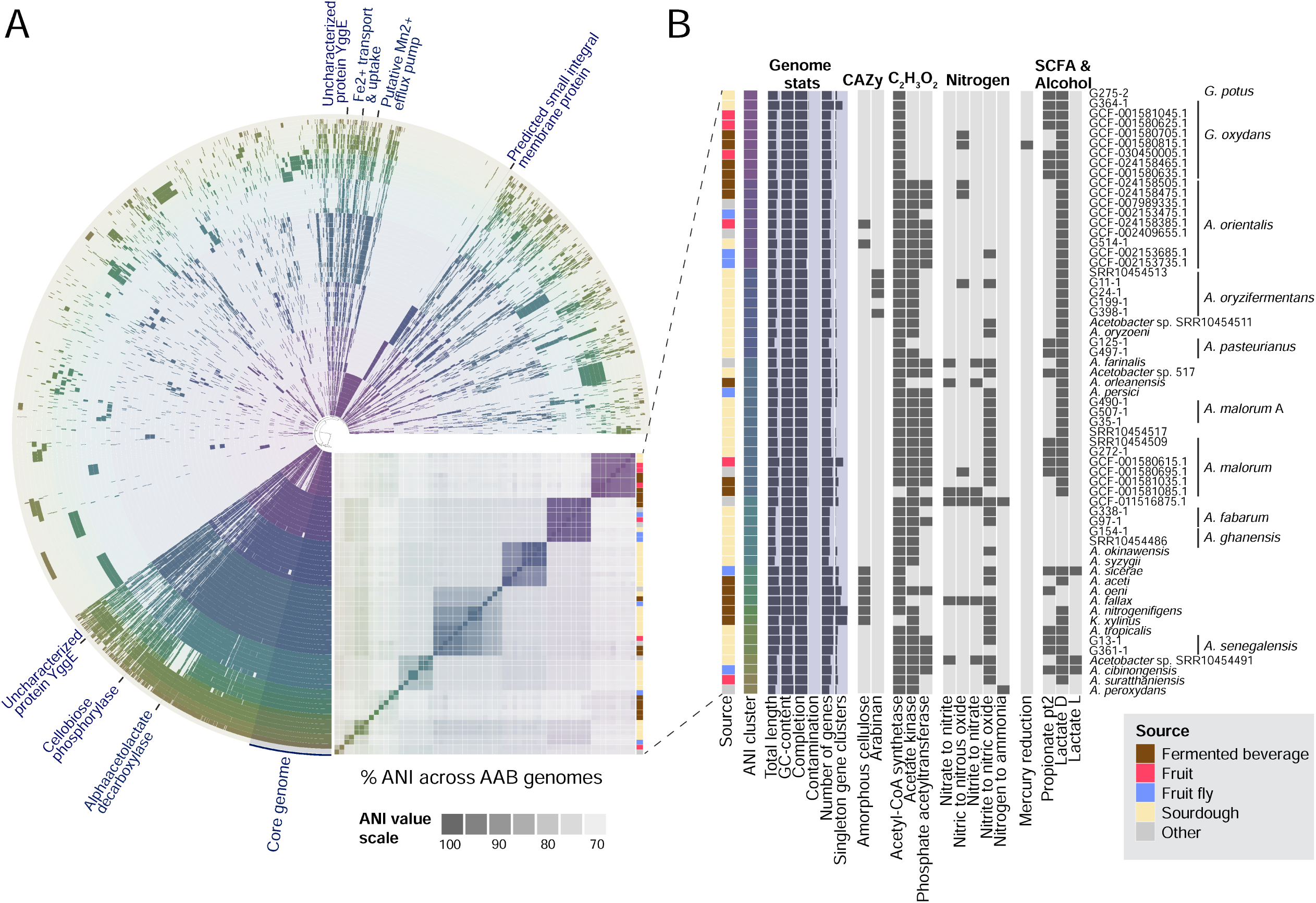
Genomic and metabolic diversity across AAB. (A) Pangenome and ANI cluster of 61 AAB genomes. Each ring represents a genome, colored by ANI cluster (**Table S5**). The core genome across AAB is highlighted, along with a subset of genes functionally enriched in sourdough genomes (for a full set, see **Table S4**). **(B)** Genome features of the 61 genomes included in the pangenome are reported by isolate source (including fermented beverage, fruit, fruit fly, sourdough, and other), ANI cluster, genome statistics (including % completion and % contamination, **Table S2**), and metabolic traits that were variable across AAB genomes, including CAZy families, acetate pathways, nitrogen metabolism, SCFA and alcohol production). Genomes are labeled by species. Where multiple genomes per species are included, Refseq ID/original name is given.

There were notable differences within and between species **(Fig. 3B**, **Table S7)** and source environment (**Table S8**) related to key metabolic traits including cellulose production and arabinan utilization. Genes for cellulose production were detected in several fermented beverage genomes within *A. aceti* and *A. nitrogenifigens*. In contrast, cellulose production was detected in only one sourdough genome, *A. orientalis* (**Fig. 3B, Table S7**). In kombucha, the source of *A. nitrogenifigens*, cellulose production is a favored trait to form the pellicle used to inoculate future batches (42, 43). In contrast, cellulose production may be a disfavored trait in sourdough due to interference with structural properties. Arabinan utilization genes were detected in four out of five *A. oryzifermentans* isolated from sourdough. Arabinan is a plant polysaccharide and the ability to break it down likely has functional consequences within the sourdough starter microbiome, which is fed by wheat flour.

### Intraspecies variation may favor persistence of AAB with key traits related to resource utilization in the sourdough environment

To determine if any genes were enriched in sourdough relative to other environments at the strain and species cluster levels (where % ANI amongst members ranged from 90-99%), we next focused our analyses on a subset of three species for which we had both sourdough and non-sourdough genome representatives: *A. malorum*/*malorum* A, *A. orientalis*, and *G. oxydans/potus*. We note that both *A. malorum* A and *G. potus* were recently reclassified as distinct species from *A. malorum* and *G. oxydans*, respectively. We first performed pangenomic analyses on nine genomes within each of the three species clusters. The total gene clusters ranged from 3,998 (*A. orientalis*) to 6,207 (*A. malorum/malorum* A) (**Fig. 4A**, **Table S6, Fig. S3**). On average, species core genomes comprised ∼40% of gene clusters whereas the other 60% of gene clusters were accessory. There was no clustering of overall genome content by either isolation source or geographic location within these species (**Fig. 3A**). This could be because many of the AAB in sourdough starter microbiomes are successful generalists adapted to a wide range of environments rather than one specific niche. They may have a nomadic lifestyle similar to some species of LAB such as *L. plantarum* (44, 45) and the AAB species *A. senegalensis* (46). Alternatively, the inclusion of more AAB genomes from sourdough and other environments may reveal signatures of adaptation that are environment specific. Despite an overall lack of clustering by geography or isolation source, we found potential gene functions that were enriched in sourdough across multiple strains. GH13 alpha-amylase, involved in starch cleavage, was enriched in sourdough with presence in 45% of strains relative to 28% of strains not isolated from sourdough (*P* < 0.01; **Fig. 4C, Table S9**). Starch is a major component of wheat, so this could confer competitive advantage to AAB in sourdough versus other environments (47, 48).

**Fig. 4:**
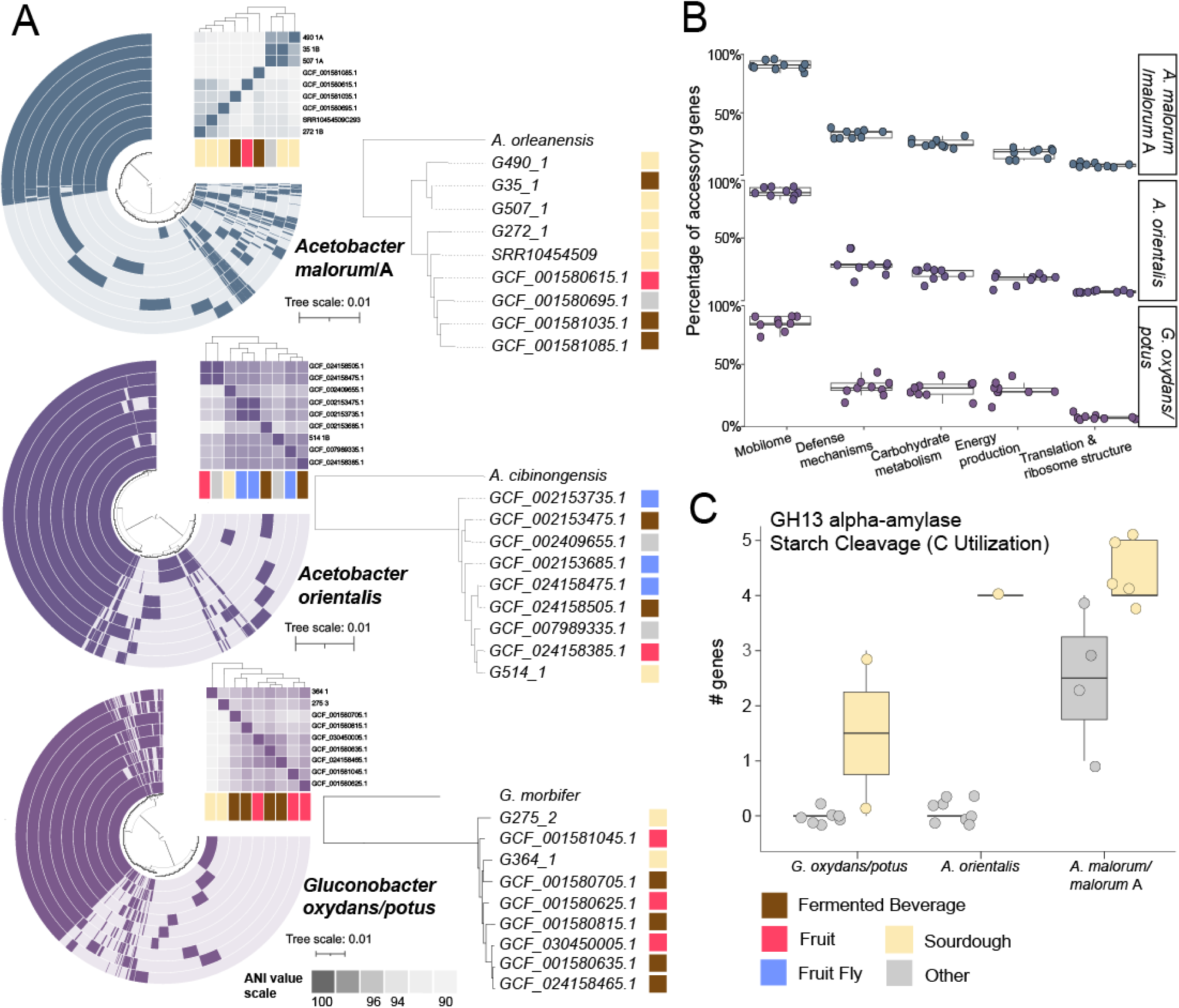
Diversity within AAB species. (A) Strain pangenomes with ANI heat map (**Table S5**) and corresponding genome trees (SpeciesTree) of nine strains each of *A. malorum/*A, *A. orientalis*, and *G. oxydans/potus*. Pangenomes and trees are annotated by the isolation source. **(B)** Percentage of genes in the accessory genome within a subset of functional categories (mobilome, defense mechanisms, carbohydrate metabolism, energy metabolism, and translation) across the three species highlighting strain diversity. **(C)** Boxplot highlights GH13 alpha-amylase (starch cleavage) which was found to be enriched in sourdough starter versus other environments in multiple strains of the same species.

Although limited by the availability of AAB genomes from other environments, our analyses start to shed light on the metabolic traits that may favor persistence of AAB in the sourdough environment.

While gene clusters of unknown function were much more frequent in accessory genomes, functionally-annotated accessory genes largely fell into categories that may have relevance to success in sourdough. On average, 61% of genes were annotated with a known function (COG database) in accessory versus 87% in core (**Table S6**). Although expected, this underscores the difficulty in determining the functional or ecological significance of gene content variation. Mobilome genes, related to transposons and prophages, made up the largest percentage of accessory genes (**Fig. 4B; Fig. S2; Table S10)**. *A. pasteurianus* have been known to have a high level of genetic variability particularly linked to the mobilome, with 9% of genes encoding transposases (49). Twenty two carbohydrate-active enzymes (CAZy) that may be relevant to success in sourdough were found in plasmids and carried by prophages in the genomes we recovered from sourdough, including the GH13 alpha-amylase significantly associated with sourdough genomes (**Fig. S4**). Notably, 24% of genes related to carbohydrate utilization were detected in accessory genomes, underscoring the variability in microbial traits related to nutrient acquisition. Accessory carbohydrate utilization genes included 5-carboxyvanillate decarboxylase LigW, involved in lignin degradation, and alpha-galactosidase (GalA). The high intraspecies variation in resource utilization genes may impact the establishment success and persistence of the AAB strain within the sourdough microbiome.

### Synthetic common garden experiments reveal distinct species and strain level effects of acetic acid bacteria on overall microbiome function

To understand the consequences of acetic acid bacteria on the assembly and emergent function of sourdough starter microbiomes, we constructed replicate synthetic sourdough starter communities, varying only the AAB strain across treatments (**Fig. 5A)**. We selected ten AAB strains that represented the breadth of phylogenetic diversity observed and also included intraspecies diversity within the *A. malorum* cluster. We qualitatively observed morphological differences in size and color across the ten strains (**Fig. S5**). For example, *Acetobacter sp. nov. 517* and *G. potus* colonies were smaller whereas *A. malorum* and *A. malorum* A were the largest. This could be due to differences in starting concentrations, nutritional requirements, or growth rates. Most colonies were white to pink in color but *A. fabarum* had yellow colonies (**Fig. S5**). *A. orientalis* was observed to produce a biofilm, likely bacterial cellulose, or another exopolysaccharide. Cellulose is produced by many AAB including those isolated from kombucha (50, 51) but amorphous colonies due to cellulose production were not commonly observed in the AAB strains isolated from sourdough.

**Fig. 5:**
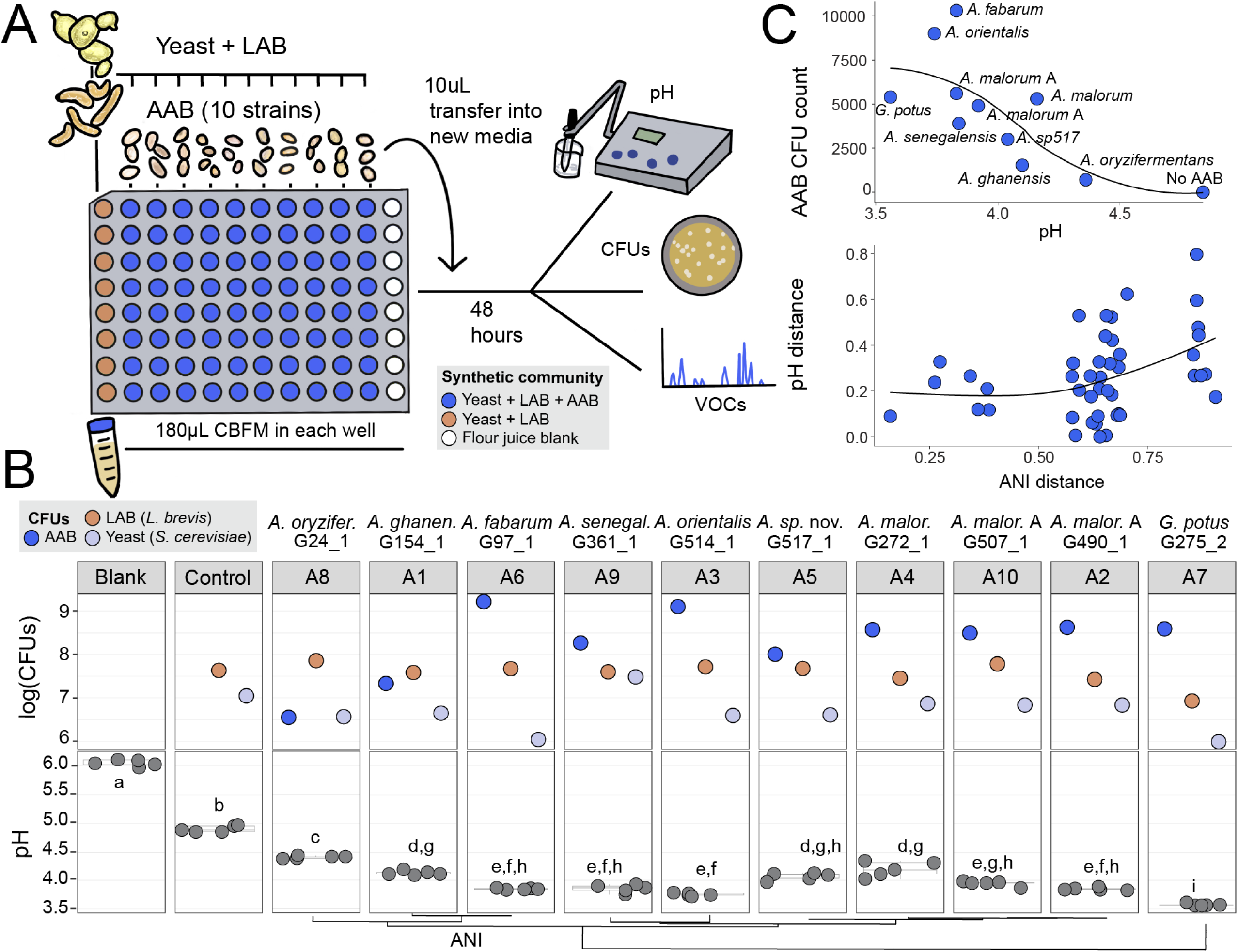
Sourdough starter acidification is determined by differences in genus, species, and strain-level variation in acetic acid bacteria. (A) Experimental design of synthetic communities. Ten isolates of AAB were selected as treatment groups and added to a background starter community of a yeast (*S. cerevisiae*) and lactic acid bacteria (*L. brevis*). We also included a yeast + LAB only control and CBFM blank where no microbes were added. All isolates were added in 5 ul at a total density of 20,000 CFUs. **(B)** Total abundance of each member measured via CFUs and plotted with log10 (top); acidification measured via pH from the liquid CBFM and reported at the end of 4 days of incubation and transfers (n = 5 replicate synthetic communities; dots represent individual observations and box plots summarize the distribution). **(C)** Plots of the relationships between AAB CFU count and emergent microbiome pH (top) and ANI versus pH distance (bottom).

For our synthetic experiments, we selected an LAB (*L. brevis*) and yeast (*S. cerevisiae*) as the base community, isolated from wild starter S_129 (resulting in ‘SynCom129’) varying only the AAB strain added. We also included a media-only blank (all synthetic communities were grown in cereal-based fermentation medium (CBFM; (14)), a liquid-based cereal fermentation media) and a yeast and LAB (YL)-only control. This design resulted in 12 distinct treatments (**Table S12**). After incubating for 48 hours, transfer to fresh media, and incubation for another 48 hours (four days total), we measured the resultant community structure and emergent functions across treatments. We assessed the persistence of all starting members and emergent functions of the resultant starter including acidification (pH) and VOC profiles. We selected a second background community of *L. brevis* and *S. cerevisiae* (SynCom361) isolated from a second wild sourdough (S_361) to confirm the trends we observed with pH and persistence analyses.

In both synthetic communities, all ten AAB strains persisted across replicates. SynCom129 was isolated from a starter without AAB, suggesting that AAB are likely able to establish and persist in a range of starters. While yeast and LAB CFU counts stayed fairly consistent, AAB CFU counts varied greatly by strain, indicating potential differential success of strains in the synthetic communities. Higher AAB CFUs were significantly correlated with lower pH (-0.83, *P* = 0.002) (**Fig. 5C, Table S14**). pH was also correlated with AAB ANI (*P* = 0.055; **Fig. 5C, Table S14)**. These results suggest that AAB density and differences in genome content may play a role in modulating starter acidification. The yeast and LAB persisted in every condition in SynCom129, but in SynCom361, the yeast sometimes failed to persist. This could be due to a mismatch of nutritional requirements but also may reflect inhibition by the AAB. We observed yeast dropout in one condition where yeast appeared glued in EPS from *A. orientalis* (**Fig. S6**).

All AAB treatments acidified the starter environment, and the extent was strain-specific (**Fig. S7**). In SynCom129, in comparison to the YL-only control, AAB overall decreased the pH of the sourdough microbiome by 18.5%. On day 4, all communities with AAB had significantly lower pH from YL-only and CBFM blank (*P* < 0.0001; **Fig. 5B, Table S14**). The average pH of communities with AAB on day 4 was 3.94 (n = 50, SD = 0.23), while the average for YL-only was 4.84 (n = 5, SD = 0.05). Significant acidification was also observed in SynCom361on day 4 between AAB and non-AAB communities (*P* < 0.0001), despite yeast dropout in some replicates (**Fig. S7**). This is consistent with studies from other fermented foods that have shown that AAB lower the pH of their environment (6, 7). The release of acetic acid and corresponding decrease in pH may inhibit the growth of other species and aid in AAB persistence (52). While pH was lowered in all AAB treatments, there were significant differences between species, including between strains of closely related species. *A. oryzifermentans* (average 4.36, SD = 0.02) and *G. potus* (average 3.56, SD = 0.02) were significantly different from all other AAB strains. The two *A. malorum* A strains did not significantly differ from each other, while the strain of *A. malorum* differed from only *A. malorum* A 490. This finding underscores the importance of species and strain-level diversification on emergent function.

Sourdough VOCs significantly differed by starter community members. We analyzed the VOCs from seven different communities including six AAB treatments and a yeast-LAB only control (**Table S13**). VOCs in the YL community significantly differed from all AAB communities (*P* = 0.008, **Table S14**), and the AAB communities formed three clusters (A4 alone, A2/A6/A7, and A8/A10, **Fig. 6A**) with distinct VOC profiles (*P* = 0.001) (**Fig. 6B; Table S14**). Clusters did not associate with phylogeny. Notably, despite 97% similarity in ANI, *A. malorum* A 507 and *A. malorum* A 490 were in different clusters. A variety of compounds differed between the clusters (**Table S13; Fig. 6B**). Stearic acid (*P* = 0.02; AAB-associated clusters) and palmitic acid (*P* = 0.02; higher the no-AAB controls and cluster A4) have both been previously detected in ferments including Tarhana (a Turkish wheat and yogurt-based fermented food (53)). We also detected compounds previously linked to flavor formation in sourdough including decanal, which is associated with vinegar and a sweet, citrusy flavor (54), and L-arabitol, which is associated with sweet flavor (55). Both were more abundant in AAB+ samples although not significantly (**Table S13**). Our results suggest that strain-level variation in AAB members can impact sensory properties of the resulting community.

**Fig. 6:**
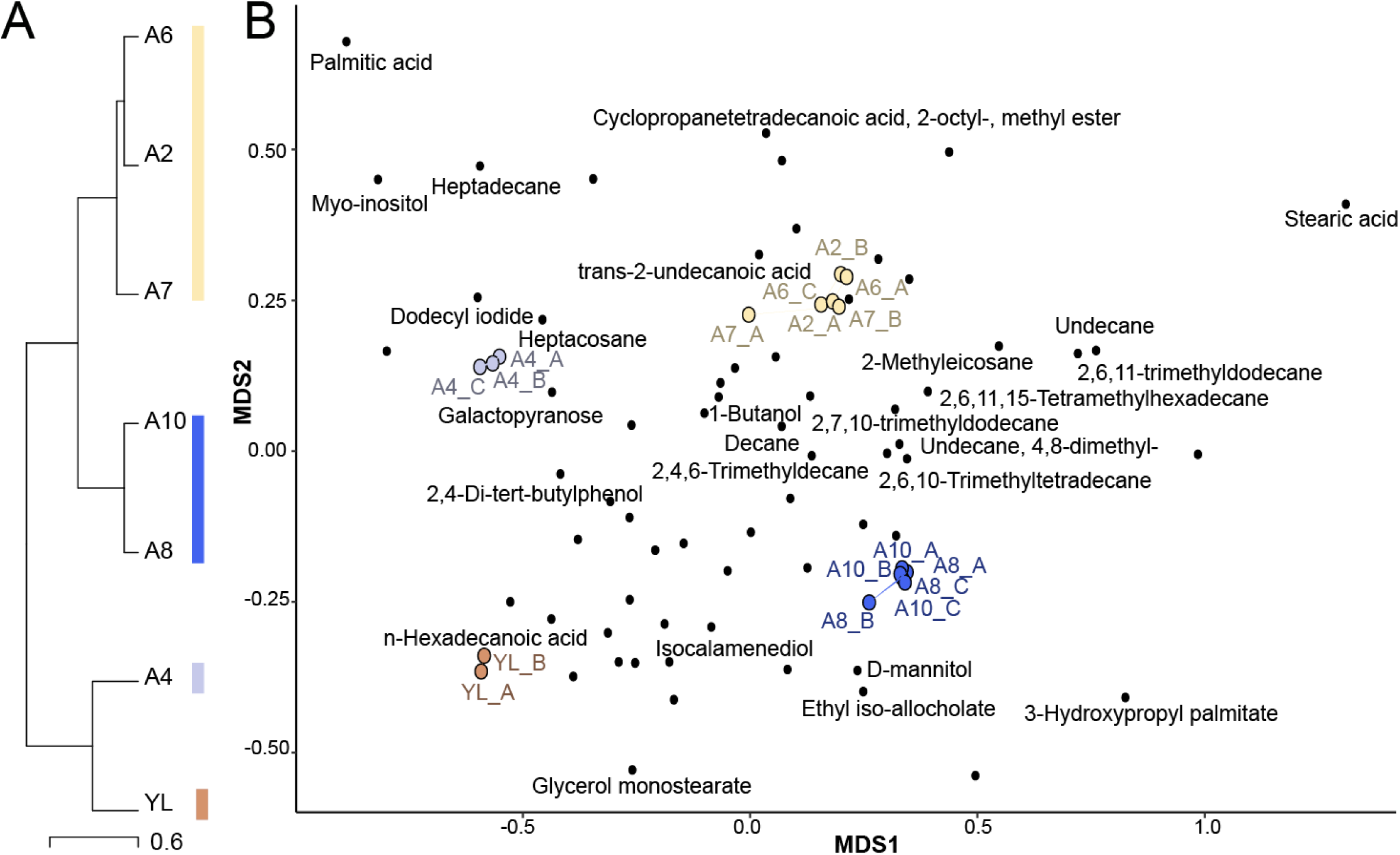
Sourdough VOCs are associated with microbial community type. (A) Dendrogram of VOC hierarchical clustering resulting in four sample clusters. **(B)** Superimposed NMDS plots of liquid sourdough communities and VOCs by community. The four main clusters have been colored. VOCs have been labeled when significantly different between clusters.

## Conclusion

Our study represents the first comprehensive genomic and ecological characterization of AAB isolated from sourdough starters and demonstrates the utility of synthetic consortia to determine functional consequences of specific microbiome members. Our genomic analyses highlight the diversity in AAB genome content and suggest that sourdough starter AAB may be enriched in specific gene functions that facilitate success in the sourdough starter environment. Our synthetic experiments demonstrate that phylogenetically and metabolically diverse AAB can persist in common sourdough microbiomes. We also determine the impact of AAB species and strain-level diversity on emergent functions including the acidification and volatile profiles of sourdough. Notably, we found that intraspecies variation in *A. malorum* resulted in significantly different community VOC profiles of synthetic starters.

These findings have direct relevance to industrial and home-bakers considering flavor profiles and other features of sourdough bread. A clear next step is to extend these results and assess how AAB may impact functional consequences in baked sourdough bread. Similarly, our synthetic experiments were performed using a *S. cerevisae* and *L. brevis* background community. Although this pairing represents one of the most dominant sourdough LAB and yeast combinations, it will be important to determine if AAB have similar functional consequences in other common sourdough types such as *F. sanfranciscensis* and *Kazachstania humilis*. An exciting avenue for future research is using tools such as interaction screens, RNA-seq, and metabolomics to shed light on what molecular-level mechanisms underlie the observed community-wide functional shifts.

## Methods

### Isolation and sequencing of acetic acid bacteria strains

We leveraged previously sequenced 16S rRNA gene amplicon data of 500 sourdough starters (14) to selectively target starter samples that had a high relative abundance of AAB and to capture a broad range of phylogenetically diverse AAB in our culturing efforts. We obtained isolates of 21 acetic acid bacteria, most of which were abundant in samples based on the amplicon data and were isolated from frozen (-80 °C) sourdough starter samples. Subsamples were plated onto selective GYCA medium (per liter, 30g glucose, 5g yeast extract, 3g peptone, 15g agar) with 10g CaCO_3_ supplemented with 25mL natamycin (21.6 mg/L) and subsamples that had amplicon hits to *Gluconobacter* were plated onto Carr medium supplemented with natamycin (21.6 mg/L) (56). To construct synthetic starter communities, we also obtained isolates of *L. brevis* (LAB) which was plated onto Lactobacilli MRS agar (Criterion) with natamycin (21.6 mg/L) and of *Saccharomyces cerevisiae* which was plated on Yeast Potato Dextrose (YPD) medium with chloramphenicol (50 mg/L). Distinct morphologies were cultured from each sample, and we used Sanger sequencing to determine general taxonomic identity with the 16s rRNA primer sets 27F/1492R for bacteria and ITS1F/ITS4R for yeast. Isolate cultures were stored in 15% glycerol at −80°C.

To sequence isolate genomes, DNA was extracted from grown colonies using ZymoBIOMICS DNA extraction kits, with bead bashing for cell lysis (Seqcenter, PA). Following extraction, sample libraries were prepped with the Illumina DNA prep kit with 10bp dual indices (IDT) and sequenced on an Illumina NovaSeq 6,000 resulting in 151 bp, paired end reads. We targeted 200 Mbp per sample (e.g. isolate). After sequencing reads were demultiplexed and adapters were trimmed using bcl-convert v4.1.5, we obtained an average of 5,676,503.62 paired-end reads per sample (ranging from 3,703,604 - 10,182,868) and an average % bp > Q30 of 91.096%.

To assemble high quality draft genomes from raw reads for each isolate in KBase (57), reads were trimmed to remove low quality bases at both ends with BBTools v38.22 (58), and low complexity reads were filtered out with PRINSEQ (59). Read quality was assessed with FastQC v0.11.0 prior to assembly with SPAdes v3.15.3. For a few genomes (N=6) the metaSPAdes assembly algorithm (60) resulted in a higher quality assembly. All genomes obtained were high quality, with ≥ 93.53% complete and < 3.23% contaminated, and the median number of contigs was 29. After assessing the genome quality of isolates with CheckM v1.0.18 (61), genomes were taxonomically classified with GTDB-Tk v1.7.0 (39, 40). We also used GTDB-Tk to assess taxonomic novelty of isolates based on reported ANI and RED values (39, 40). Genomes that did not fall within 95% ANI of existing reference genomes are considered putatively novel species. We also placed all genomes phylogenetically using KBase SpeciesTree v2.2.0 (57), which constructs species trees from a set of 49 universal core genes defined by COG gene families to determine relatedness. Assemblies were annotated with Prokka v1.14.5 (62) and re-annotated in anvi’o ((63); see comparative genomics section below) with anvi-run-hmms and anvi-run-ncbi-cogs (64).

### Inclusion of additional genomes from sourdough metagenomes and NCBI

To complement the AAB genomes that we obtained by culturing from sourdough starter samples, we also assembled genomes from 40 shotgun metagenomic samples, a subset of the 500-starter collection (14), deposited in NCBI under PRJNA589612. After quality filtering with BBDuk (65) and filtering reads that aligned to the bread wheat genome (IWGSC CS RefSeq v2.1) using bowtie2 (66), we used metaSPAdes v. 3.15.5 (60) to assemble contigs. Next, contigs were placed into genome bins with both VAMB (67) and Metabat v. 2.15 (68), and then DasTool v. 1.1.5 (69) was used to select the best bins for each sample. CheckM v. 1.2.2 (61) was used to score the bins. Next, bins from all samples were collected together and dRep v. 3.4 (70) was used to make a dereplicated set with the highest quality representatives at 95% ANI cutoff to represent species level genomes (parameters that deviated from defaults included: completeness, contamination, and coverage, which were respectively set to 50.0, 10.0, and 0.3). GTDB-tk v. 2.1.1 (39) was used to taxonomically classify the full set of dereplicated bins. We then filtered the bins based on taxonomy to only include AAB bins (N = 10 out of 30). We also built a genome tree of isolates and MAGs to identify any redundancy between genomes and MAGs. Where overlap was detected, the isolate was selected for downstream analysis and the MAG was excluded. This resulted in a final set of eight MAGs.

To contextualize our AAB sourdough genomes within the broader context of AAB genomic diversity and to assess if any gene content was enriched in sourdough environments, we also obtained publicly available AAB genomes from NCBI to include in our analyses. We found no publicly available genomes from *Acetobacter, Gluconobacter,* or *Komagataeibacter* isolated from sourdough on the JGI GOLD genomes database. For an analysis of strain diversity, we added genomes from NCBI to a set of genomes we obtained from sourdough from *A. malorum* (recently split with *A. malorum* A, ∼90% ANI)*, A. orientalis*, and *G. oxydans* (recently split with *G. potus,* ∼94% ANI) to compile a set of nine genomes within each species (19 genomes total from NCBI with at least 89% completion and less than 6.88% contamination). These three species were chosen based on the availability of multiple genomes from a variety of sources, as most AAB species have only one to a few genomes sequenced, and *A. malorum* genomes were targeted as part of the most common cluster of AAB genomes from sourdough. We also sought to include a representative set of genomes from across the tree of *Acetobacter*. We constructed a genome tree of all species of *Acetobacter* listed on NCBI Taxonomy using SpeciesTree - v2.2.0 with a reference genome from each species (**Fig. 2C**) and downloaded 12 additional *Acetobacter* genomes from across the tree that we did not recover from sourdough, along with one species from an additional genus, *K. xylinus,* as an outgroup. We targeted genomes that were at least 90% complete with < 5% contamination and only included genomes that had available source and location information. As sources often did not include exact latitude or longitude, the center point of the country or city was chosen using latlong.net.

To assess how well the genomes we recovered in this study represent the known diversity of AAB in sourdough, we leveraged our 16S rRNA gene sequencing of 500 sourdough starters. First, we assigned ASV taxonomy using a phylogeny-guided approach. We extracted full or partial 16S rRNA gene sequences from isolate, MAG, and NCBI genomes using ContEst16S (71), and aligned the sequences along with our partial 16S ASV sequences using the SINA aligner (72). After gap trimming (gap threshold = 0.2), we constructed a phylogenetic tree using RAxML-HPC2 on XSEDE (**Fig. S1)**. ASV taxonomic assignments were then made to the nearest genome representative. When an ASV was close to genomes from multiple species due to limited phylogenetic resolution, we assigned a cluster identity (such as the *A. malorum - cerevisiae* cluster).

### Comparative genomic analyses

In total, we included 61 AAB genomes in our cross-environment comparative analysis (21 isolates, 8 MAGs, and 32 from NCBI). All 61 genomes were annotated with DRAM (73) to additionally profile a suite of microbial traits related to a suite of categories including short chain fatty acid and alcohol conversions, nitrogen metabolism, and carbohydrate-active enzymes (CAZy). We display a subset of these functions that are differential across AAB (**Fig. 3B**). We also constructed a pangenome of all 61 genomes in anvi’o (minbit 0.5, mcl-inflation 6, min-occurrence 2) (74, 75) and computed ANI using pyANI (76). We annotated genomes by source (sourdough or other) and tested for functional enrichment by source and species using the command ‘anvi-compute-functional-enrichment-in-pan’ (77), a tool that finds functions that are enriched in the specified group and relative to genomes outside the group. We used the same approach to build the three species-level pangenomes for *A. malorum/malorum* A, *A. orientalis*, and *G. oxydans/potus* with corresponding ANI and functional enrichment analyses, but using mcl-inflation 10 and no min occurrence. We also identified viruses and plasmids within the genomes recovered from sourdough using geNomad and ran DRAMv on viral regions and DRAM on plasmids to annotate genes in those regions (**Fig. S4; Table S10**).

### Construction of synthetic starter communities

For synthetic sourdough starter experiments, we used a cereal-based fermentation medium (CBFM) that approximates the dough environment, as described in Landis and Oliverio et al. 2021 (14). Briefly, to make the media, whole wheat and all-purpose flour (Bob’s Red Mill) were mixed with water in a 1:1:9 ratio, centrifuged (3,000 rpm) to pellet flour particles, and then filtered through a 0.20 µm filter to remove microbial cells from the filtrate. To measure the effects of AAB on starter microbiome composition and function, we constructed two LAB and yeast communities: SynCom129, LAB *L. brevis* strain 129 plus yeast *S. cerevisiae* strain 129, and Syncom361, *L. brevis* strain 361 1A plus *S. cerevisiae* strain 361. These two species are dominant in sourdough starters and frequently found together in the same starter (14).

Additionally, this LAB/yeast pairing is commonly found in starters with AAB (the case for strains 361) and also in starters without AAB (the case for strains 129). We then selected a subset of ten AAB isolates corresponding to ten treatment groups that spanned the breadth of phylogenetic diversity we obtained: two genera (*Acetobacter* and *Gluconobacter*), eight species (*G. potus*, *A. malorum*, *A. sp.* 517, *A. orientalis*, *A. senegalensis*, *A fabarum*, *A. ghanensis*, and *A. oryzifermentans*), and three strains within *A. malorum/malorum* A. We also included a ‘no AAB’ treatment control group (only LAB/yeast) and a blank with just CBFM.

In 96-well plates, approximately 5 μl of 4000 CFUs/μL were inoculated into 180 μl of CBFM and 5 μl PBS resulting in a total volume of 200 μl (10 μl PBS was added for no AAB control and 20 μl PBS was added for blank). For each condition (ten AAB strains + one no AAB + one CBFM blank) we constructed eight replicate communities, resulting in 96 synthetic communities in total for each SynCom. Communities were incubated aerobically at room temperature on the bench and 10 μl of the resultant culture was transferred 48 hr after initial inoculation into 190 μl of fresh CBFM. At this point, wells were tested for presence/absence and pH was collected. After the cultures grew for another 48 hr (4 days in total), we harvested the experiment: three wells from each condition of the SynCom129 plate (180 μl per well) were immediately frozen at -80 °C for metabolite analyses and 100 μl from one well of each condition was subsampled to determine total abundance of each member (AAB, LAB, yeast) by serial dilution and plating on selective media at 10^-4^ and 10^-5^ to count colony forming units (CFUs). Additionally, pH was assessed *in situ* from the remaining unfrozen communities in SynCom129 and the corresponding communities in SynCom361, and one well of each condition was streaked for presence/absence (see below).

### Assessment of metabolites and acidification

All statistical tests were executed in the R environment (R Core Team). To measure the overall acidification of sourdough starter communities, the pH of each synthetic sourdough community (N = 216, excluding those that were sampled for metabolomics to avoid contamination) was taken with a pHenomenal pH meter (VWR) with a MI-410 Combination microprobe (Microelectrodes, Inc.) at the conclusion of the experiment. For our AAB synthetic community experiments, we used analyses of variance (ANOVA) to determine the effects of AAB on community pH, with AAB strain identity as the independent variable. We included a no AAB control (i.e. *L. brevis* and *S. cerevisiae* only) that we refer to as the ‘background community’ and also a CBFM only blank. We then used Tukey’s post hoc significance testing to assess the significance of pairwise differences (**Fig. 5B**). To assess if pH was correlated with the density of AAB, LAB, or yeast, we used spearman correlations (**Fig. 5C**). To determine if either the observed pH values or CFU counts were correlated with AAB phylogenetic distance, we used mantel tests (method = spearman) to correlate the pairwise distances (pH distance metric used was euclidean), with 9,999 permutations (**Fig. 5C**).

To prepare samples for GC/MS analysis, 180 μl aliquots were harvested from the final communities excluding the control and immediately frozen at -80 °C. Samples were shipped to Creative Proteomics (Shirley, NY) and upon arrival were thawed on ice and 0.1mL of each of the 21 samples was transferred to a tube with 0.3mL 80% methanol solution and 5uL ribitol (5 mg/mL). The tubes were vortexed for 60 seconds and ground for 180 seconds at 60 Hz, sonicated for 30 minutes at 4 °C, and centrifuged at 12,000 rpm and 4 °C for 10 minutes (Eppendorf).

200uL of supernatant was transferred into a new tube, dried under nitrogen, then added to 35 uL O-Methylhydroxylame solution (Merck), vortexed for 30 seconds, and incubated at 37 °C for 90 minutes. The extractives were derived with 35 uL BSTFA, reacted at 70 °C for 60 minutes, and kept at room temperature for 30 minutes until analysis. Samples were analyzed on a DB-5 column (60 m x 0.25 mm x 0.25 μm, Agilent). Detector and injector temperature were 280 °C. Oven temperature was held at 70 °C for five minutes then raised to 200 °C at a rate of 10 °C per minute, then raised to 280 °C at 5 °C per minute and held for 10 minutes. Helium was used for column carrier gas at a constant flow rate of 1 mL/min and split (20:1) injection mode was used. The temperature of the ion source was 230 °C, the electron impact mass spectra were recorded at 70 eV ionization energy, and the GC-MS analysis was carried out in 30 - 550 mass range scanning mode (Thermo Scientific ISQ 7000).

To analyze the variation of VOCs across the samples sent for GC/MS analysis, any duplicated compound reads were removed, only keeping the highest similarity matches of each compound. Compounds related to plastic components/degradation, human metabolites, or low similarity compounds were also removed as background or contamination. Then, distance matrices were calculated and ordinated of the VOCs and samples (**Fig. S6B**). Four outlier samples were removed prior to downstream analyses (YL_C, A2_C, A6_B, A7_C). To assess the impact of AAB treatment on overall community VOC profiles, we ran PERMANOVA models with treatment (AAB strain) and AAB presence (versus yeast-LAB only) as predictors. We identified four clusters of samples using hierarchical clustering (method Ward D2) and plotted the resulting dendrogram (**Fig. S6A**). We used Kruskal-Wallis tests (with FDR corrections for multiple comparisons) to assess if any VOCs were significantly enriched by AAB presence, treatment, or cluster.

## Data availability

Raw reads and assemblies for all genomes generated in this study have been deposited in the NCBI Sequence Read Archive in BioProject PRJNA1095457. Annotation and pangenome files and DRAM output have been deposited to Figshare DOI 10.6084 under (10.6084/m9.figshare.25403626, 10.6084/m9.figshare.25403620, and 10.6084/m9.figshare.25403599, respectively).

## Acknowledgements

We thank Dr. Austin Garner for his assistance in imaging the acetic acid bacteria cultures, presented in Fig. S5. This study was supported by grant #2328528/2328529 from the National Science Foundation Division of Molecular and Cellular Bioscience to A.M.O. and B.E.W.

## Supplemental Material Figure Captions

**Fig. S1: Phylogenetic tree of 16S rRNA gene sequences (N = 197) to determine ASV taxonomic assignments.** When present, full or partial 16S rRNA genes were extracted from isolate genomes, MAGs, and NCBI genomes. A phylogenetic tree was built using RAxML, and ASV taxonomic assignments were then made to the nearest genome representative (see Methods for additional details).

**Fig. S2: Full summary of core vs. accessory COG categories.** Percentage of genes in the accessory genome within all functional categories of COG annotations across the three species highlighting strain diversity (*A. malorum*/*malorum* A, *A. orientalis*, and *G. oxydans*/*potus*).

**Fig. S3: Genome by shared gene content.** Accumulation plots visualizing number of genomes included in pangenome and the resulting number of unique gene clusters for **(A)** *A. malorum/A. malorum A*, **(B)** *A. orientalis* and **(C)** *G. oxydans/G. potus.* Each point represents a unique set of genomes, and at every distinct number of genomes, all possible combinations of genomes were subsampled out of the total (N=9 for each). Curves show smoothed conditional means (loess) and gray shade indicates standard error.

**Fig. S4: CAZymes are carried on mobile elements in AAB genomes from sourdough.** Number of hits to CAZyme genes detected within sourdough AAB genomes, colored by species, and patterned by source, either on plasmids or integrated from prophages.

**Fig. S5: Variation in colony morphology of sourdough AAB.** Ten strains of acetic acid bacteria selected for synthetic starter experiments plated on GYCA with calcium carbonate, imaged with a Sony Alpha 7RIII. Scale bars represent approximately 2mm.

**Fig. S6: Yeast appear glued in exopolysaccharides from *A. orientalis*.** SynCom361 community A3 (*S. cerevisiae, L. brevis, A. orientalis*) well 8 streaked for presence/absence on day 4 onto Yeast Potato Dextrose plate with chloramphenicol (selective for yeast).

**Fig. S7: Diverse AAB species and strains all acidify the sourdough starter environment relative to the yeast and LAB-only controls and strain level differences in emergent acidification are highly conserved.** To assess acidification of the overall microbiome with and without AAB strains, we measured the pH at day 2 (after 48 hours) and day 4 (after 96 hours) of five replicate consortia for every distinct treatment. There were 204 distinct pH measurements, representing ten AAB treatments, a blank, and a YL control, with five replicates each across the two timepoints. We excluded two mistransferred wells and a subset of treatments from SynCom361 day 4 (n = 34) where yeast failed to persist. Different letters represent post hoc significant pairwise differences.

## Tables

**Table S1.** Species’ assignments of AAB ASVs recovered from 500 sourdough starters. **Table S2.** Yeast and LAB ASVs statistically enriched in AAB-dominant sourdough starters. **Table S3.** Co-occurrences of ASVs with AAB in sourdough starters.

**Table S4.** Metadata of 61 AAB genomes, including species, sources, genome stats, and genome accessions.

**Table S5.** ANI matrix of 61 AAB genomes by percent ANI between all genomes.

**Table S6.** Number of gene clusters recovered in AAB pangenomes with core and accessory distributions.

**Table S7.** Summary of microbial metabolic traits across 61 AAB genomes output from DRAM.

**Table S8.** Functional enrichments by environment (sourdough or other) across 61 AAB genomes (Anvi’o).

**Table S9.** Functional enrichments by environments (sourdough or other; Kruskal-Wallis) between AAB strains.

**Table S10.** Annotation outputs of sourdough AAB genes recovered on plasmids or from prophages.

**Table S11.** Analyses by plate and well of synthetic starter communities, including presence/absence, pH, CFU counts, and VOCs.

**Table S12.** Volatile organic compound peak areas of synthetic sourdough communities. Similarity scores match with reference ions, CAS = Chemical Abstract Service, Rt = Retention time, QC = quality control samples.

**Table S13.** VOC means and statistical differences in compound relative abundance between sample clusters.

**Table S14.** Statistical outputs relating differences in function (including pH and VOC profiles) and composition (AAB CFUs) to differences in treatments (including treatment type and differences in ANI across AAB strains).

## References

1. Qiu X, Zhang Y, Hong H. 2021. Classification of acetic acid bacteria and their acid resistant mechanism. AMB Expr 11:29.

2. Yang H, Chen T, Wang M, Zhou J, Liebl W, Barja F, Chen F. 2022. Molecular biology: Fantastic toolkits to improve knowledge and application of acetic acid bacteria. Biotechnology Advances 58:107911.

3. Landis EA, Fogarty E, Edwards JC, Popa O, Eren AM, Wolfe BE. 2022. Microbial Diversity and Interaction Specificity in Kombucha Tea Fermentations. mSystems 7:e00157–22.

4. Marsh AJ, O’Sullivan O, Hill C, Ross RP, Cotter PD. 2014. Sequence-based analysis of the bacterial and fungal compositions of multiple kombucha (tea fungus) samples. Food Microbiology 38:171–178.

5. Harrison K, Curtin C. 2021. Microbial Composition of SCOBY Starter Cultures Used by Commercial Kombucha Brewers in North America. Microorganisms 9:1060.

6. Bouchez A, De Vuyst L. 2022. Acetic Acid Bacteria in Sour Beer Production: Friend or Foe? Front Microbiol 13:957167.

7. De Roos J, De Vuyst L. 2018. Acetic acid bacteria in fermented foods and beverages. Current Opinion in Biotechnology 49:115–119.

8. Adler P, Frey LJ, Berger A, Bolten CJ, Hansen CE, Wittmann C. 2014. The Key to Acetate: Metabolic Fluxes of Acetic Acid Bacteria under Cocoa Pulp Fermentation-Simulating Conditions. Appl Environ Microbiol 80:4702–4716.

9. He Y, Xie Z, Zhang H, Liebl W, Toyama H, Chen F. 2022. Oxidative Fermentation of Acetic Acid Bacteria and Its Products. Front Microbiol 13:879246.

10. Mamlouk D, Gullo M. 2013. Acetic Acid Bacteria: Physiology and Carbon Sources Oxidation. Indian J Microbiol 53:377–384.

11. Chouaia B, Gaiarsa S, Crotti E, Comandatore F, Degli Esposti M, Ricci I, Alma A, Favia G, Bandi C, Daffonchio D. 2014. Acetic Acid Bacteria Genomes Reveal Functional Traits for Adaptation to Life in Insect Guts. Genome Biology and Evolution 6:912–920.

12. Lievens B, Hallsworth JE, Pozo MI, Belgacem ZB, Stevenson A, Willems KA, Jacquemyn H. 2015. Microbiology of sugar-rich environments: diversity, ecology and system constraints. Environmental Microbiology 17:278–298.

13. Calvert MD, Madden AA, Nichols LM, Haddad NM, Lahne J, Dunn RR, McKenney EA. 2021. A review of sourdough starters: ecology, practices, and sensory quality with applications for baking and recommendations for future research. PeerJ 9:e11389.

14. Landis EA, Oliverio AM, McKenney EA, Nichols LM, Kfoury N, Biango-Daniels M, Shell LK, Madden AA, Shapiro L, Sakunala S, Drake K, Robbat A, Booker M, Dunn RR, Fierer N, Wolfe BE. 2021. The diversity and function of sourdough starter microbiomes. eLife 10:e61644.

15. De Vuyst L, Van Kerrebroeck S, Harth H, Huys G, Daniel H-M, Weckx S. 2014. Microbial ecology of sourdough fermentations: Diverse or uniform? Food Microbiology 37:11–29.

16. Minervini F, De Angelis M, Di Cagno R, Gobbetti M. 2014. Ecological parameters influencing microbial diversity and stability of traditional sourdough. International Journal of Food Microbiology 171:136–146.

17. Gänzle M, Ripari V. 2016. Composition and function of sourdough microbiota: From ecological theory to bread quality. International Journal of Food Microbiology 239:19–25.

18. Calabrese FM, Ameur H, Nikoloudaki O, Celano G, Vacca M, Junior WJfl, Manzari C, Vertè F, Di Cagno R, Pesole G, De Angelis M, Gobbetti M. 2022. Metabolic framework of spontaneous and synthetic sourdough metacommunities to reveal microbial players responsible for resilience and performance. Microbiome 10:148.

19. Comasio A, Verce M, Van Kerrebroeck S, De Vuyst L. 2020. Diverse Microbial Composition of Sourdoughs From Different Origins. Front Microbiol 11:1212.

20. Ripari V, Gänzle MG, Berardi E. 2016. Evolution of sourdough microbiota in spontaneous sourdoughs started with different plant materials. International Journal of Food Microbiology 232:35–42.

21. Li Z, Li H, Bian K. 2016. Microbiological characterization of traditional dough fermentation starter (Jiaozi) for steamed bread making by culture-dependent and culture-independent methods. International Journal of Food Microbiology 234:9–14.

22. Lynch KM, Zannini E, Wilkinson S, Daenen L, Arendt EK. 2019. Physiology of Acetic Acid Bacteria and Their Role in Vinegar and Fermented Beverages. Comprehensive Reviews in Food Science and Food Safety 18:587–625.

23. Li H, Fu J, Hu S, Li Z, Qu J, Wu Z, Chen S. 2021. Comparison of the effects of acetic acid bacteria and lactic acid bacteria on the microbial diversity of and the functional pathways in dough as revealed by high-throughput metagenomics sequencing. International Journal of Food Microbiology 346:109168.

24. Li H, Hu S, Fu J. 2022. Effects of acetic acid bacteria in starter culture on the properties of sourdough and steamed bread. Grain & Oil Science and Technology 5:13–21.

25. Jakob F, Steger S, Vogel RF. 2012. Influence of novel fructans produced by selected acetic acid bacteria on the volume and texture of wheat breads. Eur Food Res Technol 234:493–499.

26. Van Rossum T, Ferretti P, Maistrenko OM, Bork P. 2020. Diversity within species: interpreting strains in microbiomes. Nat Rev Microbiol 18:491–506.

27. Wolff R, Shoemaker W, Garud N. 2023. Ecological Stability Emerges at the Level of Strains in the Human Gut Microbiome. mBio 14:e02502–22.

28. Kalan LR, Meisel JS, Loesche MA, Horwinski J, Soaita I, Chen X, Uberoi A, Gardner SE, Grice EA. 2019. Strain- and Species-Level Variation in the Microbiome of Diabetic Wounds Is Associated with Clinical Outcomes and Therapeutic Efficacy. Cell Host & Microbe 25:641–655.e5.

29. Ellegaard KM, Engel P. 2019. Genomic diversity landscape of the honey bee gut microbiota. Nat Commun 10:446.

30. Viver T, Conrad RE, Rodriguez-R LM, Ramírez AS, Venter SN, Rocha-Cárdenas J, Llabrés M, Amann R, Konstantinidis KT, Rossello-Mora R. 2024. Towards estimating the number of strains that make up a natural bacterial population. Nat Commun 15:544.

31. You L, Jin H, Kwok L-Y, Lv R, Zhao Z, Bilige M, Sun Z, Liu W, Zhang H. 2023. Intraspecific microdiversity and ecological drivers of lactic acid bacteria in naturally fermented milk ecosystem. Science Bulletin 68:2405–2417.

32. Koch H, Germscheid N, Freese HM, Noriega-Ortega B, Lücking D, Berger M, Qiu G, Marzinelli EM, Campbell AH, Steinberg PD, Overmann J, Dittmar T, Simon M, Wietz M. 2020. Genomic, metabolic and phenotypic variability shapes ecological differentiation and intraspecies interactions of Alteromonas macleodii. Sci Rep 10:809.

33. Niccum BA, Kastman EK, Kfoury N, Robbat A, Wolfe BE. 2020. Strain-Level Diversity Impacts Cheese Rind Microbiome Assembly and Function. mSystems 5:e00149–20.

34. Zhou W, Spoto M, Hardy R, Guan C, Fleming E, Larson PJ, Brown JS, Oh J. 2020. Host-Specific Evolutionary and Transmission Dynamics Shape the Functional Diversification of Staphylococcus epidermidis in Human Skin. Cell 180:454–470.e18.

35. Conwill A, Kuan AC, Damerla R, Poret AJ, Baker JS, Tripp AD, Alm EJ, Lieberman TD. 2022. Anatomy promotes neutral coexistence of strains in the human skin microbiome. Cell Host & Microbe 30:171–182.e7.

36. Arrieta-Echeverri MC, Fernandez GJ, Duarte-Riveros A, Correa-Álvarez J, Bardales JA, Villanueva-Mejía DF, Sierra-Zapata L. 2023. Multi-omics characterization of the microbial populations and chemical space composition of a water kefir fermentation. Front Mol Biosci 10:1223863.

37. Sindi A, Badsha MdB, Ünlü G. 2020. Bacterial Populations in International Artisanal Kefirs. Microorganisms 8:1318.

38. Perpetuini G, Tittarelli F, Battistelli N, Suzzi G, Tofalo R. 2020. Contribution of Pichia manshurica strains to aroma profile of organic wines. Eur Food Res Technol 246:1405–1417.

39. Chaumeil P-A, Mussig AJ, Hugenholtz P, Parks DH. 2020. GTDB-Tk: a toolkit to classify genomes with the Genome Taxonomy Database. Bioinformatics 36:1925–1927.

40. Chaumeil P-A, Mussig AJ, Hugenholtz P, Parks DH. 2022. GTDB-Tk v2: memory friendly classification with the genome taxonomy database. Bioinformatics 38:5315–5316.

41. Wolfe BE, Button JE, Santarelli M, Dutton RJ. 2014. Cheese Rind Communities Provide Tractable Systems for In Situ and In Vitro Studies of Microbial Diversity. Cell 158:422–433.

42. Tran T, Grandvalet C, Winckler P, Verdier F, Martin A, Alexandre H, Tourdot-Maréchal R. 2021. Shedding Light on the Formation and Structure of Kombucha Biofilm Using Two-Photon Fluorescence Microscopy. Front Microbiol 12:725379.

43. Dutta D, Gachhui R. 2006. Novel nitrogen-fixing Acetobacter nitrogenifigens sp. nov., isolated from Kombucha tea. International Journal of Systematic and Evolutionary Microbiology 56:1899–1903.

44. Martino ME, Bayjanov JR, Caffrey BE, Wels M, Joncour P, Hughes S, Gillet B, Kleerebezem M, van Hijum SAFT, Leulier F. 2016. Nomadic lifestyle of *Lactobacillus plantarum* revealed by comparative genomics of 54 strains isolated from different habitats. Environmental Microbiology 18:4974–4989.

45. Cen S, Yin R, Mao B, Zhao J, Zhang H, Zhai Q, Chen W. 2020. Comparative genomics shows niche-specific variations of Lactobacillus plantarum strains isolated from human, Drosophila melanogaster, vegetable and dairy sources. Food Bioscience 35:100581.

46. Almeida OGG, Gimenez MP, De Martinis ECP. 2022. Comparative pangenomic analyses and biotechnological potential of cocoa-related Acetobacter senegalensis strains. Antonie van Leeuwenhoek 115:111–123.

47. Liu Y, Yu J, Li F, Peng H, Zhang X, Xiao Y, He C. 2017. Crystal structure of a raw-starch-degrading bacterial α-amylase belonging to subfamily 37 of the glycoside hydrolase family GH13. Sci Rep 7:44067.

48. Katina K, Salmenkallio-Marttila M, Partanen R, Forssell P, Autio K. 2006. Effects of sourdough and enzymes on staling of high-fibre wheat bread. LWT - Food Science and Technology 39:479–491.

49. Azuma Y, Hosoyama A, Matsutani M, Furuya N, Horikawa H, Harada T, Hirakawa H, Kuhara S, Matsushita K, Fujita N, Shirai M. 2009. Whole-genome analyses reveal genetic instability of Acetobacter pasteurianus. Nucleic Acids Research 37:5768–5783.

50. Chen S-Q, Lopez-Sanchez P, Wang D, Mikkelsen D, Gidley MJ. 2018. Mechanical properties of bacterial cellulose synthesised by diverse strains of the genus Komagataeibacter. Food Hydrocolloids 81:87–95.

51. Dayal MS, Goswami N, Sahai A, Jain V, Mathur G, Mathur A. 2013. Effect of media components on cell growth and bacterial cellulose production from Acetobacter aceti MTCC 2623. Carbohydrate Polymers 94:12–16.

52. Xia M, Zhang X, Xiao Y, Sheng Q, Tu L, Chen F, Yan Y, Zheng Y, Wang M. 2022. Interaction of acetic acid bacteria and lactic acid bacteria in multispecies solid-state fermentation of traditional Chinese cereal vinegar. Front Microbiol 13:964855.

53. Ovando-Martinez M, Daglioglu O, Guner KG, Gecgel U, Simsek S. 2014. Analysis of the Fatty Acids and Phenolic Compounds in a Cereal-Based Fermented Food (Tarhana). FNS 05:1177–1184.

54. Kong H, Kim SH, Jeong W-S, Kim S-Y, Yeo S-H. 2023. Microbiome and Volatile Metabolic Profile of Acetic Acid Fermentation Using Multiple Starters for Traditional Grain Vinegar. Fermentation 9:423.

55. Du P, Yuan H, Chen Y, Zhou H, Zhang Y, Huang M, Jiangfang Y, Su R, Chen Q, Lai J, Guan L, Ding Y, Hu H, Luo J. 2023. Identification of Key Aromatic Compounds in Basil (Ocimum L.) Using Sensory Evaluation, Metabolomics and Volatilomics Analysis. Metabolites 13:85.

56. Subbiahdoss G, Osmen S, Reimhult E. 2022. Cellulosic biofilm formation of Komagataeibacter in kombucha at oil-water interfaces. Biofilm 4:100071.

57. Arkin AP, Cottingham RW, Henry CS, Harris NL, Stevens RL, Maslov S, Dehal P, Ware D, Perez F, Canon S, Sneddon MW, Henderson ML, Riehl WJ, Murphy-Olson D, Chan SY, Kamimura RT, Kumari S, Drake MM, Brettin TS, Glass EM, Chivian D, Gunter D, Weston DJ, Allen BH, Baumohl J, Best AA, Bowen B, Brenner SE, Bun CC, Chandonia J-M, Chia J-M, Colasanti R, Conrad N, Davis JJ, Davison BH, DeJongh M, Devoid S, Dietrich E, Dubchak I, Edirisinghe JN, Fang G, Faria JP, Frybarger PM, Gerlach W, Gerstein M, Greiner A, Gurtowski J, Haun HL, He F, Jain R, Joachimiak MP, Keegan KP, Kondo S, Kumar V, Land ML, Meyer F, Mills M, Novichkov PS, Oh T, Olsen GJ, Olson R, Parrello B, Pasternak S, Pearson E, Poon SS, Price GA, Ramakrishnan S, Ranjan P, Ronald PC, Schatz MC, Seaver SMD, Shukla M, Sutormin RA, Syed MH, Thomason J, Tintle NL, Wang D, Xia F, Yoo H, Yoo S, Yu D. 2018. KBase: The United States Department of Energy Systems Biology Knowledgebase. Nat Biotechnol 36:566–569.

58. BBMap – Bushnell B. – sourceforge.net/projects/bbmap/.

59. Schmieder R, Edwards R. 2011. Quality control and preprocessing of metagenomic datasets. Bioinformatics 27:863–864.

60. Nurk S, Meleshko D, Korobeynikov A, Pevzner PA. 2017. metaSPAdes: a new versatile metagenomic assembler. Genome Res 27:824–834.

61. Parks DH, Imelfort M, Skennerton CT, Hugenholtz P, Tyson GW. 2015. CheckM: assessing the quality of microbial genomes recovered from isolates, single cells, and metagenomes. Genome Res 25:1043–1055.

62. Seemann T. 2014. Prokka: rapid prokaryotic genome annotation. Bioinformatics 30:2068–2069.

63. Eren AM, Kiefl E, Shaiber A, Veseli I, Miller SE, Schechter MS, Fink I, Pan JN, Yousef M, Fogarty EC, Trigodet F, Watson AR, Esen ÖC, Moore RM, Clayssen Q, Lee MD, Kivenson V, Graham ED, Merrill BD, Karkman A, Blankenberg D, Eppley JM, Sjödin A, Scott JJ, Vázquez-Campos X, McKay LJ, McDaniel EA, Stevens SLR, Anderson RE, Fuessel J, Fernandez-Guerra A, Maignien L, Delmont TO, Willis AD. 2020. Community-led, integrated, reproducible multi-omics with anvi’o. Nat Microbiol 6:3–6.

64. Hyatt D, Chen G-L, LoCascio PF, Land ML, Larimer FW, Hauser LJ. 2010. Prodigal: prokaryotic gene recognition and translation initiation site identification. BMC Bioinformatics 11:119.

65. Bushnell B. 2014. BBTools software package.

66. Langmead B, Salzberg SL. 2012. Fast gapped-read alignment with Bowtie 2. Nat Methods 9:357–359.

67. Nissen JN, Johansen J, Allesøe RL, Sønderby CK, Armenteros JJA, Grønbech CH, Jensen LJ, Nielsen HB, Petersen TN, Winther O, Rasmussen S. 2021. Improved metagenome binning and assembly using deep variational autoencoders. Nat Biotechnol 39:555–560.

68. Kang DD, Li F, Kirton E, Thomas A, Egan R, An H, Wang Z. 2019. MetaBAT 2: an adaptive binning algorithm for robust and efficient genome reconstruction from metagenome assemblies. PeerJ 7:e7359.

69. Sieber CMK, Probst AJ, Sharrar A, Thomas BC, Hess M, Tringe SG, Banfield JF. 2018. Recovery of genomes from metagenomes via a dereplication, aggregation and scoring strategy. Nat Microbiol 3:836–843.

70. Olm MR, Brown CT, Brooks B, Banfield JF. 2017. dRep: a tool for fast and accurate genomic comparisons that enables improved genome recovery from metagenomes through de-replication. The ISME Journal 11:2864–2868.

71. Lee I, Chalita M, Ha S-M, Na S-I, Yoon S-H, Chun J. 2017. ContEst16S: an algorithm that identifies contaminated prokaryotic genomes using 16S RNA gene sequences. International Journal of Systematic and Evolutionary Microbiology 67:2053–2057.

72. Pruesse E, Peplies J, Glöckner FO. 2012. SINA: Accurate high-throughput multiple sequence alignment of ribosomal RNA genes. Bioinformatics 28:1823–1829.

73. Shaffer M, Borton MA, McGivern BB, Zayed AA, La Rosa SL, Solden LM, Liu P, Narrowe AB, Rodríguez-Ramos J, Bolduc B, Gazitúa MC, Daly RA, Smith GJ, Vik DR, Pope PB, Sullivan MB, Roux S, Wrighton KC. 2020. DRAM for distilling microbial metabolism to automate the curation of microbiome function. Nucleic Acids Research 48:8883–8900.

74. Edgar RC. 2004. MUSCLE: multiple sequence alignment with high accuracy and high throughput. Nucleic Acids Research 32:1792–1797.

75. van Dongen S, Abreu-Goodger C. 2012. Using MCL to Extract Clusters from Networks, p. 281–295. *In* van Helden, J, Toussaint, A, Thieffry, D (eds.), Bacterial Molecular Networks. Springer New York, New York, NY.

76. Pritchard L, Glover RH, Humphris S, Elphinstone JG, Toth IK. 2016. Genomics and taxonomy in diagnostics for food security: soft-rotting enterobacterial plant pathogens. Anal Methods 8:12–24.

77. Shaiber A, Willis AD, Delmont TO, Roux S, Chen L-X, Schmid AC, Yousef M, Watson AR, Lolans K, Esen ÖC, Lee STM, Downey N, Morrison HG, Dewhirst FE, Mark Welch JL, Eren AM. 2020. Functional and genetic markers of niche partitioning among enigmatic members of the human oral microbiome. Genome Biol 21:292.

